# Structures of the *Staphylococcus aureus* ribosome inhibited by fusidic acid and fusidic acid cyclopentane

**DOI:** 10.1101/2023.11.17.567578

**Authors:** Adrián González-López, Daniel S. D. Larsson, Ravi Kiran Koripella, Brett N. Cain, Martin Garcia Chavez, Paul J. Hergenrother, Suparna Sanyal, Maria Selmer

## Abstract

The antibiotic fusidic acid (FA) is used to treat *Staphylococcus aureus* infections. It inhibits protein synthesis by binding to elongation factor G (EF-G) and preventing its release from the ribosome after translocation. While FA is only effective against gram-positive bacteria, the available structures of FA-inhibited complexes are from gram-negative model organisms. To fill this knowledge gap, we solved cryo-EM structures of the *S. aureus* ribosome in complex with mRNA, tRNA, EF-G and FA to 2.5 Å resolution and the corresponding complex structures with the recently developed FA derivative FA-cyclopentane (FA-CP) to 2.0 Å resolution. With both FA variants, the majority of the ribosomal particles are observed in chimeric hybrid state and only a minor population in post-translocational state. As expected, FA binds in a pocket between domains I, II and III of EF-G and the sarcin-ricin loop of 23S rRNA. FA-CP binds in an identical position, but its cyclopentane moiety provides additional contacts to EF-G and 23S rRNA, suggesting that its improved resistance profile towards mutations in EF-G is due to higher-affinity binding. These high-resolution structures reveal new details about the *S. aureus* ribosome, including confirmation of many rRNA modifications, and provide an optimal starting point for future structure-based drug discovery on an important clinical drug target.

## INTRODUCTION

Fusidic acid (FA) is a bacteriostatic antibiotic that inhibits bacterial protein synthesis. It is effective mainly against gram-positive bacteria and used routinely against topical *Staphylococcus aureus* infections since the 1960s (1). However, due to resistance development (2, 3), FA is usually recommended at high doses and in combination with other antibiotics (4, 5). The development of new FA derivatives has been largely unsuccessful, as modifications of the drug usually come at the cost of potency (6, 7).

FA targets elongation factor G (EF-G), the GTPase factor that catalyzes translocation of mRNA and tRNAs on the ribosome (8). FA inhibits the release of EF-G from the ribosome and thereby stalls translation (9, 10). Translocation requires two major conformational changes of the ribosome: intersubunit rotation and rotation of the head of the small ribosomal subunit (SSU), known as swiveling (11, 12). FA also locks EF-G in the recycling step of translation (13, 14), where EF-G together with ribosome recycling factor dissociates the tRNAs and mRNA from the ribosome (15). The binding site of FA is a pocket between domains I-III of EF-G and the sarcin-ricin loop (SRL) of 23S rRNA, which is formed after GTP hydrolysis. FA binding seems to stabilize an intermediate between the GTP and GDP conformation of EF-G, where switch I is in GDP conformation but switch II remains in GTP conformation, which disables the necessary conformational changes for EF-G release (16). High-resolution structural information for the FA-inhibited ribosome was first obtained from X-ray crystallography with EF-G and ribosomes from *Thermus thermophilus* (16–20), and more recently from cryo-EM studies with components from *Escherichia coli* (21). However, all these structures are from gram-negative bacteria, most of which are inherently resistant to FA (22). From the actual FA target pathogen, *S. aureus*, there are only structures of EF-G without the ribosome (23).

The FA-locked structures have exhibited two different ribosomal states. One is the post-translocational state (POST) (19), where the ribosome has completed translocation, and tRNAs are bound in the canonical P and E sites. The other is an intermediate so-called chimeric hybrid state (CHI) of translocation, in which the head of the small ribosomal subunit (SSU) remains swiveled and tRNAs make A- and P-site interactions with the mRNA codon and the SSU head, but P- and E-site interactions with the 30S body and the 50S (12, 18). In both states, EF-G is in close contact with 23S rRNA, 16S rRNA, uL6, uS12, mRNA, and P-site tRNA.

There are 3 main types of FA resistance observed in *Staphylococcus sp*. The *fusA* mutants have mutations in the drug target, EF-G (2, 24). FusB-type resistance is caused by a *fusB*-encoded resistance protein (with homologous variants FusC, FusD, and FusF) that binds to EF-G and causes resistance without direct interaction with the antibiotic (25, 26). The *fusE* variants carry truncations or loss-of-function mutations in the *rplF* gene, encoding ribosomal protein uL6, which interacts with EF-G (3).

Here, we set out to determine high-resolution cryo-EM structures of the clinical FA drug target, the *S. aureus* 70S ribosome where EF-G is locked with FA or a recently developed FA derivative. These structures allowed comparison with ribosome-EF-G-FA structures from other bacteria, to re-analyze known *fusA* resistance mutations and to refine a high-quality model that can be used for the design of new FA derivatives or analogs, as well as other drugs targeting the *S. aureus* ribosome.

## RESULTS AND DISCUSSION

### Structure of the *S. aureus* ribosome

We assembled a minimal high-occupancy FA-locked *S. aureus* ribosome complex for structure determination. *S. aureus* 70S ribosomes were mixed with a short synthetic mRNA, *E. coli* tRNA^fMet^, and *S. aureus* EF-G in presence of FA and GTP and imaged by cryogenic electron microscopy (cryo-EM). Image processing using cryoSPARC (27) produced two different ribosome reconstructions in complex with EF-G bound to FA: A post-translocational state (POST) at 3.1 Å resolution, and a chimeric hybrid pe/E state (CHI) at 2.5 Å (Figure 1A).

**Figure 1.**
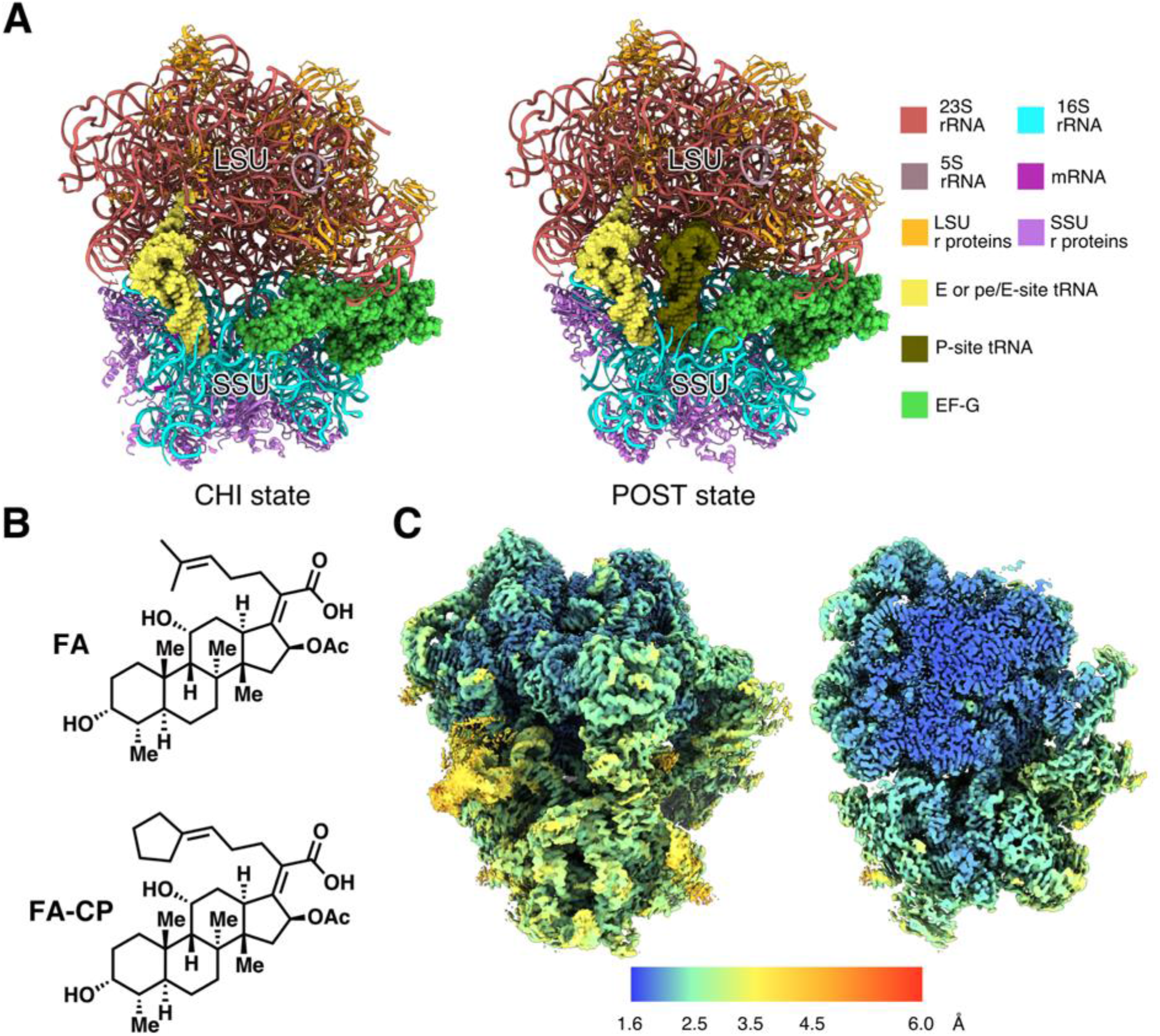
Structures of *S. aureus* ribosomes in complex with mRNA, tRNA, EF-G, and FA/FA-CP. (A) Composition of the different complexes. Large subunit (LSU) and small subunit (SSU) are indicated. (B) Chemical structure of FA and FA-CP. (C) Local resolution map over the surface and a central slice on the highest resolution map (2.0 Å FA-CP CHI state).

We then applied the same methodology to the recently developed FA derivative fusidic acid cyclopentane (FA-CP, Figure 1B) (28), which has equivalent potency against *S. aureus* as FA, but a better resistance profile. This resulted in a 2.0-Å resolution CHI state reconstruction and a 2.4-Å POST state.

The reconstructed maps are very similar, the main differences between the CHI and POST states are the head swivel of the SSU, and the resulting changes in interactions with the tRNAs and EF-G. In all complexes, the ribosomal A-site is occupied by domain IV of EF-G. The POST state contains two tRNAs in the P- and E sites, whereas the CHI state only has one tRNA bound in pe/E state (Figure 1A). The pe/E tRNA makes P-site interactions with the codon on the mRNA and the SSU head (16S nucleotides 1349-1352 and 1238-1241), but E-site interactions with the body of 16S (nucleotides 700-704), and with the 50S subunit. The mRNA Shine-Dalgarno sequence is base paired with the anti-Shine-Dalgarno 1546C-1551U of 16S rRNA in both CHI structures. In the POST structure, however, both the mRNA and the 16S rRNA are disordered in this region. The CHI state is the most abundant in our sample, and only a minor population is in POST state. In the FA data, 60 % of the ribosomes are bound to EF-G, and 93 % of those are in CHI state. In the FA-CP data, 72 % of the 70S particles contain EF-G, out of which 88% are in CHI state.

The map quality of FA-CP CHI complex was excellent with a majority of the large subunit reaching 1.6 Å (Figure 1C). The high resolution allowed the refinement of a high-quality model of the *S. aureus* 70S ribosome (Table 1), identification of many rRNA modifications, and correction of modeling errors in previous lower-resolution structures (*e.g.* a D159-A160 cis-peptide bond in uL3, Figure S1). This model was subsequently refined into the FA-CP POST and FA-CHI maps. Due to map anisotropy and lower quality, no modelling of the FA POST state was done.

**Table 1.**
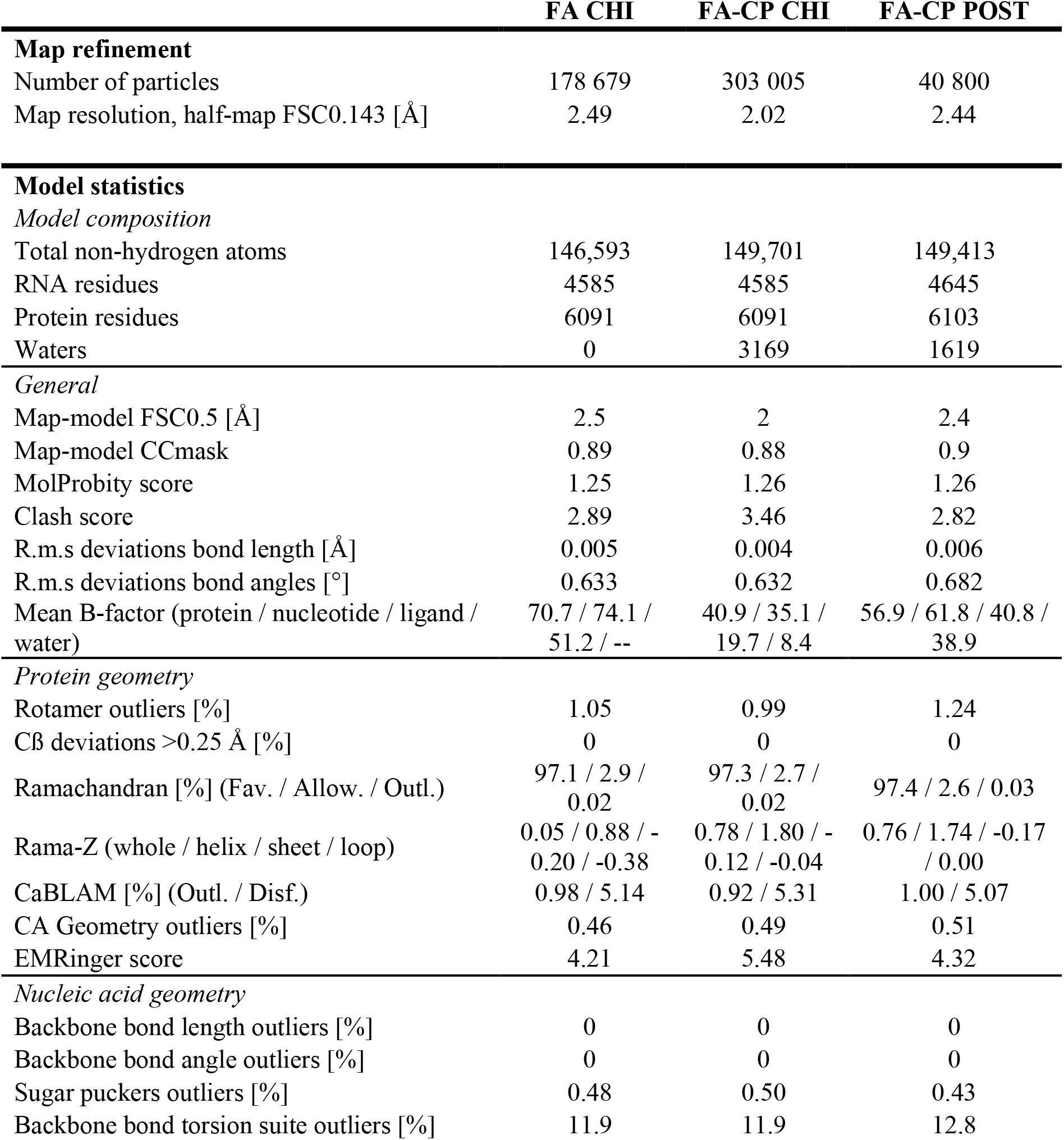
Cryo-EM refinement parameters and model validation.

|  | FA CHI | FA-CP CHI | FA-CP POST |
| --- | --- | --- | --- |
| **Map refinement** |  |  |  |
| Number of particles | 178 679 | 303 005 | 40 800 |
| Map resolution, half-map FSC0.143 [Å] | 2.49 | 2.02 | 2.44 |
| **Model statistics** |  |  |  |
| *Model composition* |  |  |  |
| Total non-hydrogen atoms | 146,593 | 149,701 | 149,413 |
| RNA residues | 4585 | 4585 | 4645 |
| Protein residues | 6091 | 6091 | 6103 |
| Waters | 0 | 3169 | 1619 |
| *General* |  |  |  |
| Map-model FSC0.5 [Å] | 2.5 | 2 | 2.4 |
| Map-model CCmask | 0.89 | 0.88 | 0.9 |
| MolProbity score | 1.25 | 1.26 | 1.26 |
| Clash score | 2.89 | 3.46 | 2.82 |
| R.m.s deviations bond length [Å] | 0.005 | 0.004 | 0.006 |
| R.m.s deviations bond angles [°] | 0.633 | 0.632 | 0.682 |
| Mean B-factor (protein / nucleotide / ligand / water) | 70.7 / 74.1 / 51.2 / -- | 40.9 / 35.1 / 19.7 / 8.4 | 56.9 / 61.8 / 40.8 / 38.9 |
| *Protein geometry* |  |  |  |
| Rotamer outliers [%] | 1.05 | 0.99 | 1.24 |
| Cβ deviations >0.25 Å [%] | 0 | 0 | 0 |
| Ramachandran [%] (Fav. / Allow. / Outl.) | 97.1 / 2.9 / 0.02 | 97.3 / 2.7 / 0.02 | 97.4 / 2.6 / 0.03 |
| Rama-Z (whole / helix / sheet / loop) | 0.05 / 0.88 / -0.20 / -0.38 | 0.78 / 1.80 / -0.12 / -0.04 | 0.76 / 1.74 / -0.17 / 0.00 |
| CaBLAM [%] (Outl. / Disf.) | 0.98 / 5.14 | 0.92 / 5.31 | 1.00 / 5.07 |
| CA Geometry outliers [%] | 0.46 | 0.49 | 0.51 |
| EMRinger score | 4.21 | 5.48 | 4.32 |
| *Nucleic acid geometry* |  |  |  |
| Backbone bond length outliers [%] | 0 | 0 | 0 |
| Backbone bond angle outliers [%] | 0 | 0 | 0 |
| Sugar puckers outliers [%] | 0.48 | 0.50 | 0.43 |
| Backbone bond torsion suite outliers [%] | 11.9 | 11.9 | 12.8 |

### *S. aureus* EF-G on the ribosome

The binding and overall conformation of the FA-bound *S. aureus* EF-G on the ribosome agrees with previous cryo-EM and X-ray crystallography structures of FA-locked complexes from gram-negative bacteria (16–19, 21). Overall, our POST structure is most similar to the *T. thermophilus* structure in the same state (PDBID: 4WQF, TtPOST), whereas the CHI structure is more similar to CHI state structures that have both ap/P and pe/E tRNAs, from *E. coli* (PDBID: 7N2C, EcCHI) and *T. thermophilus* (PDBID: 4W29, TtCHI). Other CHI-state structures that only have one tRNA in the pe/E site (*e.g.* PDB ID: 4V9L) show a different degree of head swivel.

The conformation of EF-G is very similar in the CHI and POST states (RMSD: 0.382 Å over 666 Cα atoms of EF-G). The largest difference is found in a loop at the tip of domain IV (residues 498-505), due to additional interactions with the mRNA and P-site tRNA in the POST structure.

EF-G interacts with both 23S and 16S rRNA, as well as with ribosomal proteins uL6 and uS12 (Figure 2A). The main contacts with 23S rRNA are between domains I and V of EF-G (residues H18, H85, S660 and Q663) and the SRL proximal to the FA binding site (Figure 2A). Domain IV of EF-G interacts with 23S and 16S rRNA at the intersubunit interface, where R498 contacts helix 69 of 23S rRNA and helix 44 of 16S rRNA (Figure S2). In the *E. coli* structure, the equivalent residue (K507) does not reach the corresponding interface. In the CHI structures, we observe an additional interaction, not observed in previous structures, between S586 in EF-G and C1941 (C1914 in *E. coli*), in the hairpin loop of helix 69 of 23S rRNA at intersubunit bridge B2a (Figure S2). C1941 is observed in two conformations, it either interacts with EF-G or stacks with helix 69. The previous nucleotide, A1940 (A1913 in *E. coli*), can also adopt different conformations depending on the state of the ribosome. In our structures, it stacks with A1503 (A1492 in *E. coli*) of 16S rRNA when EF-G contacts its backbone. In absence of EF-G, it instead forms a base interaction with A-site tRNA (*e.g.* in PDB 7K00 (29)).

**Figure 2.**
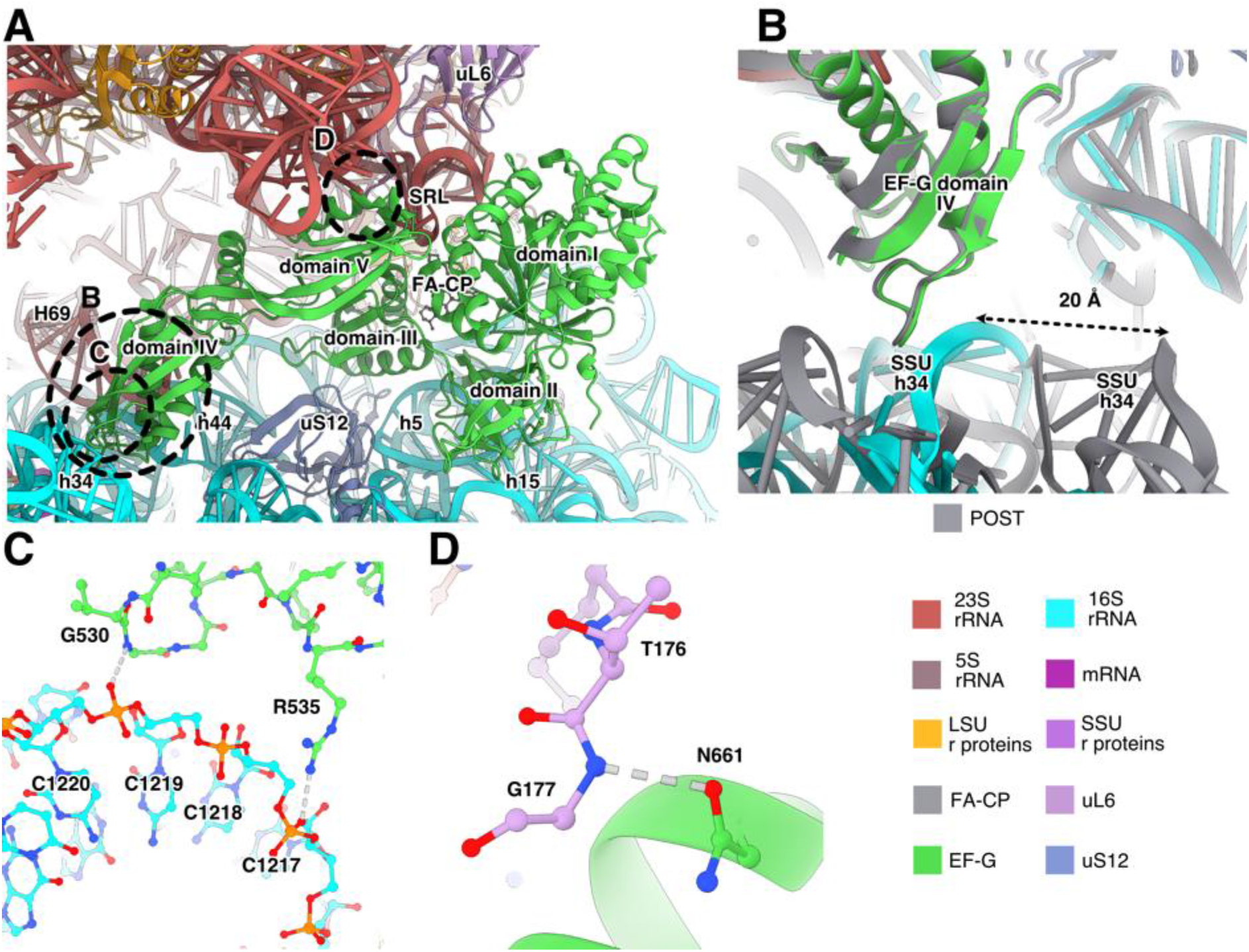
Interactions of EF-G with the *S. aureus* ribosome. (A) Binding site of EF-G in the ribosomal CHI state. Interacting helices from 23S rRNA are indicated with “H” and helices from 16S rRNA with “h”. (B) Differences between the SSU contacts of EF-G in POST (gray) and CHI state. The SSU head swivel brings 16S rRNA helix 34 in contact with EF-G. (C) Interactions of EF-G with helix 34 in the CHI state. (D) Interaction between EF-G and uL6 shown for the CHI state.

Domain II of EF-G binds to 16S rRNA helix 5 (residue K324) and helix 15 (residues Y340, R349, R351, R354). In EcCHI, the contact through R349 is not present, since the corresponding sidechain (R357 in *E. coli*) points in the opposite direction and is far from the RNA. Residue 340 is a threonine in EcCHI (T348) that does not reach the RNA. In the CHI structure, EF-G makes additional contacts with helix 34 in the head region of 16S rRNA (Figure 2B,C). This results in a 10% larger hidden surface area of EF-G compared to the POST state, suggesting that higher stability causes its higher abundance. In fact, in the recent time-resolved cryo-EM studies of EF-G translocation in *E. coli* (30, 31), EF-G was only present in pre-translocation and chimeric states. The ribosomal POST conformation was only observed after EF-G dissociation, suggesting that EF-G naturally leaves the ribosome before the POST conformation is reached. Furthermore, in cryo-EM structures of FA-inhibited *E. coli* ribosomes, only the CHI state was observed (21). In our data, however, we do observe POST state, indicating that FA inhibition allows time for the SSU head to back-swivel in presence of EF-G.

Differently from TtPOST and EcCHI, EF-G contacts uL6 through a polar interaction between N661 and the backbone of T176 (Figure 2D). This can explain why truncations of the C-terminus of uL6 (*fusE* mutations) provide low-level resistance to FA, since this will weaken the interactions of EF-G with the ribosome and facilitate its release in presence of FA.

EF-G also interacts with uS12 through residues D421, T441 and E443 in domain III (Figure 2A). In TtPOST, the interactions are similar, but residue 443 is a proline that cannot form the equivalent interaction. The interaction with uS12 in EcCHI is slightly different, *e.g.* the interaction of T441 is replaced by the equivalent residue of *S. aureus* G446 (N454).

### Binding site of FA and FA-CP

FA and FA-CP bind next to GDP and magnesium in a pocket between domains I-III of EF-G and the SRL of 23S (Figure 3A-B, Figure S3), exposed to the solvent at the ribosomal subunit interface. The main interactions of EF-G with FA are formed by switch II (residues 80-90) in domain I and helix 9 (residues 455-470) in domain III. Before GTP hydrolysis, switch I (residues 39-63) occupies this site to interact with the gamma phosphate of GTP, impeding FA binding. After GTP hydrolysis, switch I becomes disordered and switch II changes conformation. FA locks switch II in the GTP conformation, stabilizing EF-G’s interactions with the ribosome (*e.g.* H85 with 23S) and inhibiting its release (Figure S4).

**Figure 3.**
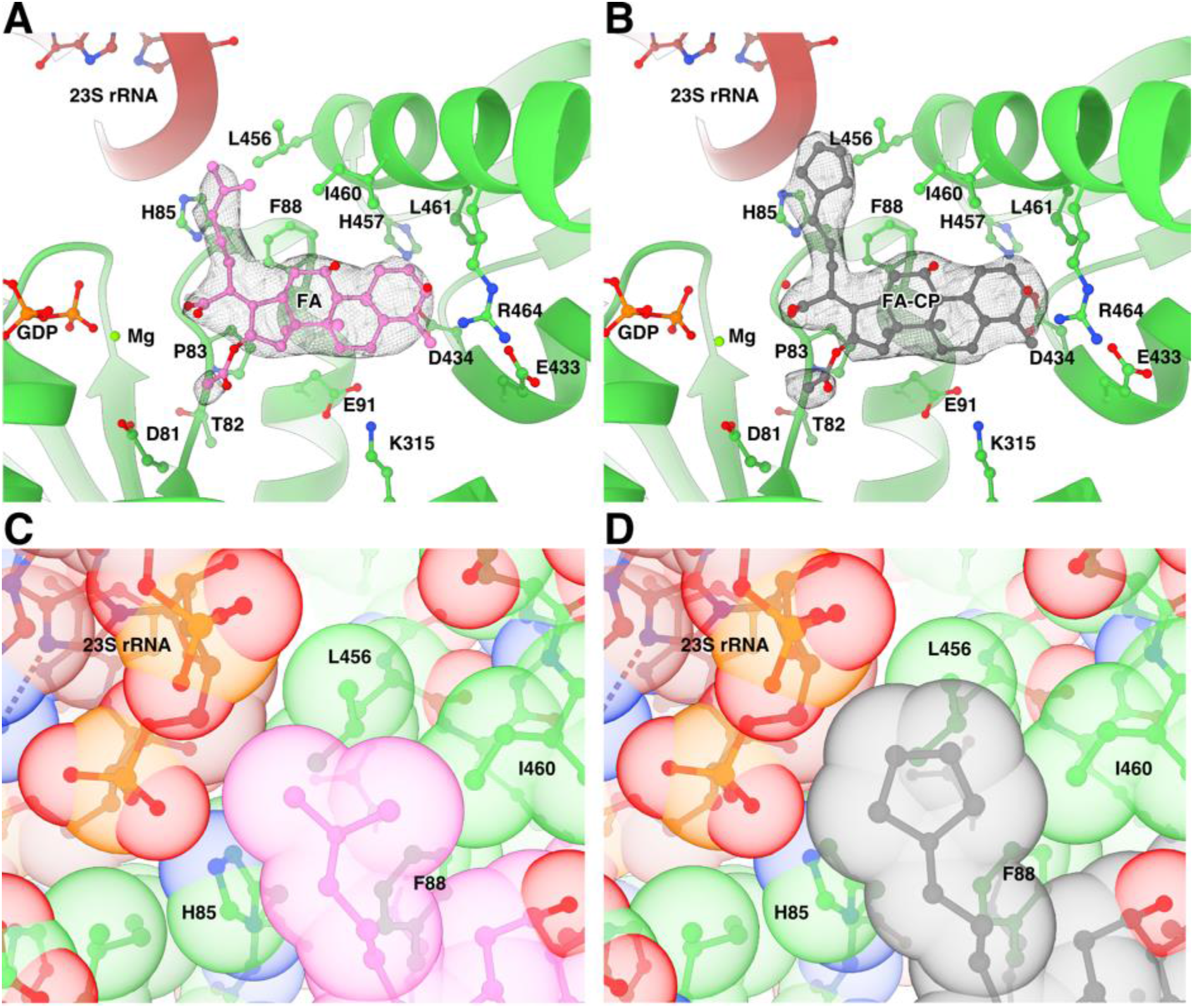
Binding site of FA and FA-CP. (A-B) Segmented local filtered maps around FA (A) or FA-CP (B) in CHI state. Side chains within 4.5 Å of FA or FA-CP are shown as sticks. (C-D) Close-up of the packing of the dimethyl alkene side chain of FA (C) and the cyclopentane variant in FA-CP (D) within the binding pocket, showed with transparent ball representation at van der Waals radii.

Our maps clearly show the extra density for the carbons of the cyclopentane moiety of FA-CP (Figure 3A–B, Figure S3), forming tighter interactions with H85, L456 and the SRL (Figure 3C-D). The interactions formed by the additional atoms in FA-CP presumably lead to increased affinity and further stabilization of the FA-bound conformation of EF-G. In line with previous structure-activity relationships, this region of the binding pocket appears to be most amenable to structure modifications as other relatively lipophilic moieties such as cyclohexane and tetrahydropyran rings can be placed in the same location with no detriment to antimicrobial activity in *S. aureus* (28). Future projects aiming for improved activity against *S. aureus* may be successful in identifying higher affinity derivatives by thoroughly probing this area with relatively non-polar functional groups.

In comparison with previously published structures, FA binds in a similar manner as in the EcCHI structure, with minor differences in the detailed sidechain conformations, particularly for I460 (Figure 4A). On the other hand, in TtPOST, residues L456 and I460 have different rotamers, switch II has a slightly different orientation with H85 farther from interaction with 23S, and helix 9 from domain III is tilted, resulting in a 2 Å movement of the Cα of R464 (Figure 4B).

**Figure 4.**
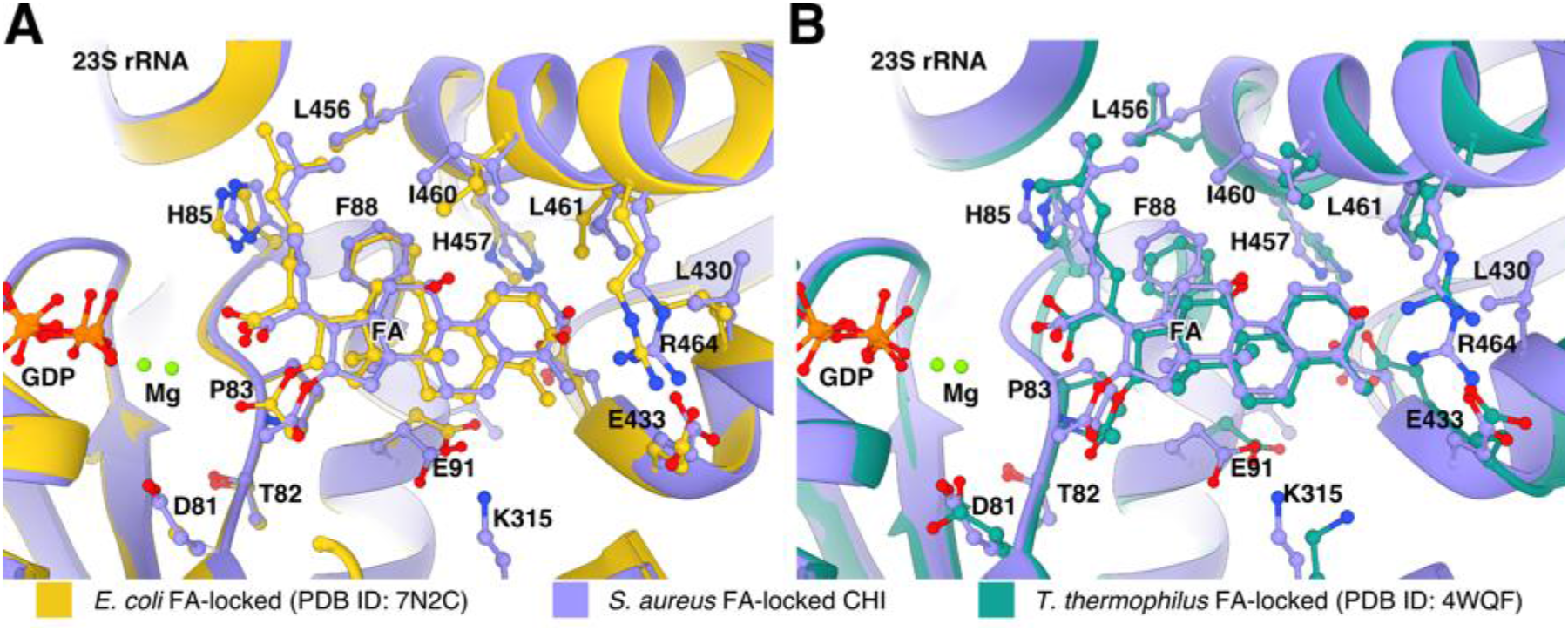
Comparison of the FA binding site in *S. aureus* complexes with previously available structures. Structures were aligned based on EF-G and side chains within 4.5 Å of FA are shown as sticks. (A) Overlay of the FA binding site in the *S. aureus* CHI state and EcCHI (PDB ID: 7N2C). (B) Overlay of the FA binding site in the *S. aureus* CHI state and TtPOST (PDB ID: 4WQF).

### *fusA* mutants

Structures of FA-locked ribosome complexes have demonstrated that FA-inhibition requires drug binding to a certain EF-G conformation with specific interactions with the ribosome. FA resistance mutations in EF-G (*fusA*-type) from clinical isolates and lab studies (3, 32–34) have been found to affect all of these aspects, and have thus been classified into four groups according to their expected mechanism of rescue: effects on FA binding, ribosome-EF-G interactions, EF-G conformation or EF-G stability (23). Based on our high-resolution structures, we analyze seven different sites of mutation where FA-CP gives lower minimum inhibitory concentrations than FA (28). Around the FA binding site, the strong F88L mutation and the milder R464A would weaken direct EF-G-FA interactions (Figure 5). The higher affinity of FA-CP may partially compensate for the absence of these interactions. The potent resistance mutations D434N and H457L/N/Y and the milder T436I would break the hydrogen-bond network between D434, T436 and H457 (at 4.3 Å distance of F88 in domain I), which bring together two helices at the drug-facing surface of domain III. Additional mutations in domain III are connected to this network by hydrophobic interaction directly: V90I; via M453: P406L (tested with FA-CP), V407F, G451V and G452C/SV; or via P435: T385N in domain II (tested with FA-CP) (Figure 5). We thus confirm that interactions in the core of domain III are important for FA potency, directly and through interactions with domains I and III. We also conclude that tighter binding of FA-CP seems to lead to increased stability of the FA-locked conformation; allowing FA-CP to partially compensate for mutation-induced non-optimal interactions in this core. It remains to be tested whether FA-CP also shows increased potency against the milder resistance mutations in domain V of EF-G.

**Figure 5.**
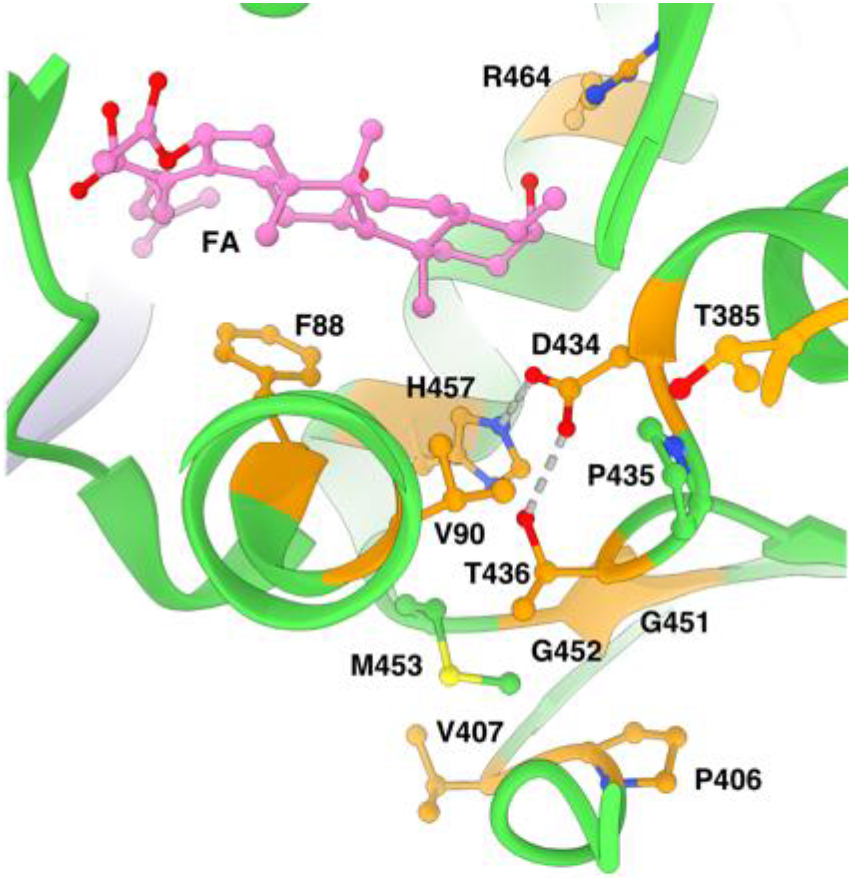
Network of interactions in EF-G around the FA binding site. Sites of *fusA* resistance mutations are shown in orange.

### *S. aureus* ribosomal RNA modifications

Based on the analysis of the 2.0 Å cryo-EM map and the comparison with *E. coli* rRNA modifications, 14 rRNA modifications could be identified (Figure 6) including two novel modifications of 23S rRNA and three novel modifications of 16S rRNA (35). The previously reported 2’-O-methylcytidine in position C1947 of 23S rRNA is not present in our structure.

**Figure 6.**
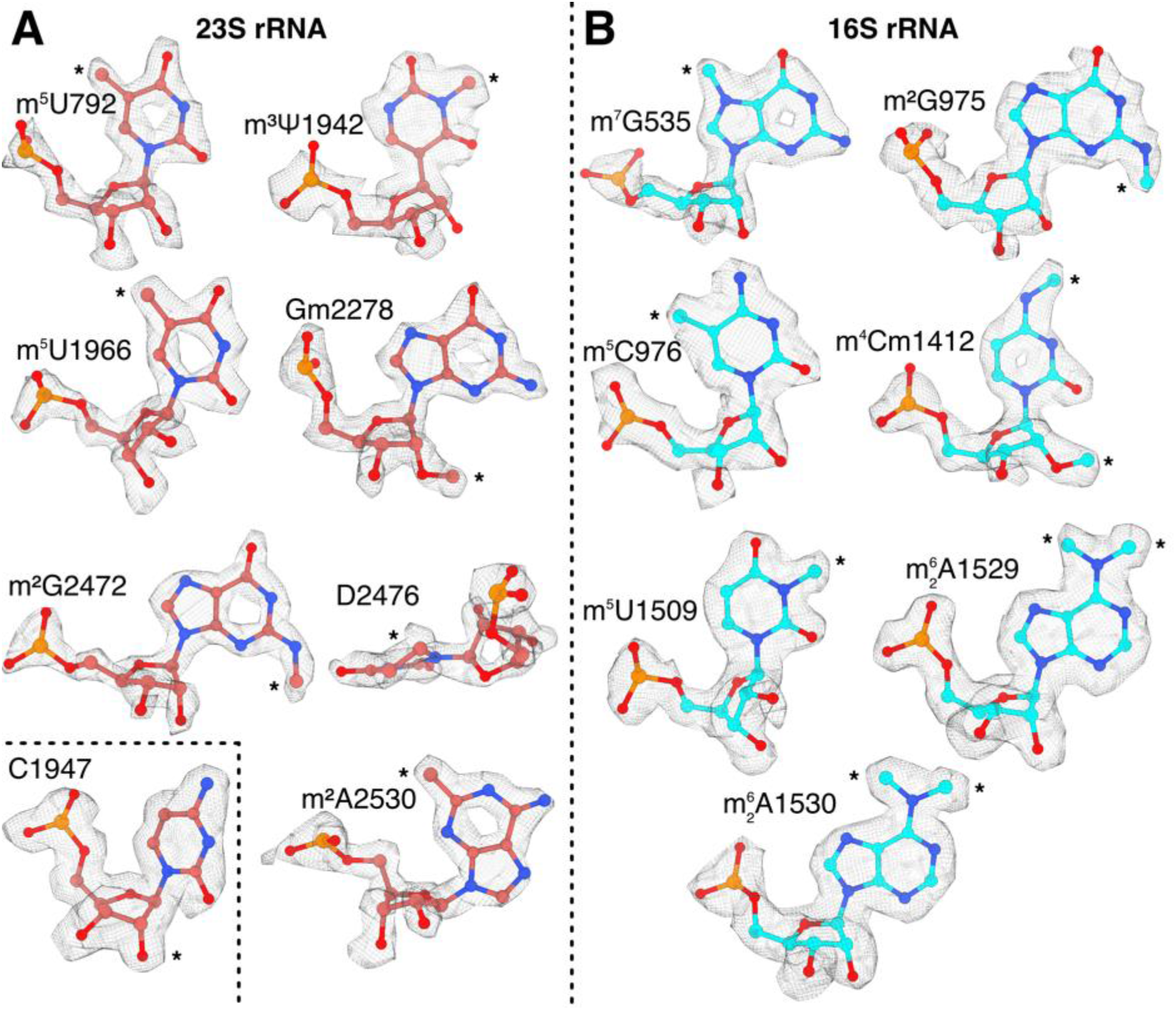
Identification of post-transcriptionally modified rRNA nucleotides in local filtered maps of the FA-CP CHI complex of the *S. aureus* ribosome. Asterisks indicate sites of modification. (A) Identified rRNA modifications in 23S rRNA. The inset shows that C1947 lacks 2’O methylation. (B) rRNA modifications in 16S rRNA.

For 23S rRNA, we identified a 5-methyluridine in position U792 (helix 35, *E. coli* U747), where the methyl moiety packs against the ribose of C2639. The corresponding modification is made by RlmC in *E. coli*. A BLAST (36) search against *S. aureus* proteins only gave hits classified as homologs of RlmD, another uridine methylase that modifies U1939. We also observed C1966 (equivalent position in *S. aureus*) to be methylated. In other gram-positive bacteria, such as *Bacillus subtilis*, both of these modifications are performed by the dual-specificity enzyme RlmCD (37). Our observations suggest that this also occurs in *S. aureus*, as earlier proposed without observation of the m5 modification of U792.

Additionally, we found clear density for a non-planar base at position 2476, located at the peptidyl-transfer center, which suggests the presence of 5,6-dihydrouridine (*E. coli* D2449). We also found density for a methylation of N3 of at position 1942, located in helix 69 of 23S. In *E. coli*, this is an N3-methylpseudouridine. There are homologues of RluD and RlmH in *S. aureus*, corroborating that the same modifications are present in *S. aureus*.

For 16S rRNA, we found a N2-methylguanosine at position G975 and 5-methylcytidine at C976, located at the loop of helix 31 (same modification in *E. coli* G966 and C967), part of the P-site. The methyl group in G975 stacks against K131 of uS9, which is interacting directly with the P-site tRNA next to the anticodon loop.

Lastly, we also found a 5-methyluridine at position U1509, in helix 44 (same modification in *E. coli* 1498). This modification is also located on the P-site, next to the mRNA codon-anticodon interaction, where also C1412 is modified to N4,O2’-dimethylcytidine. A summary of all the rRNA modifications that were confidently identified from the density can be found in Table 2.

**Table 2.** *S. aureus* rRNA modifications identified in the cryo-EM map.

| Position<br>*S. aureus* | Position<br>*E. coli* | Modification | *E. coli* enzyme | Probable *S. aureus*<br>enzyme (UniProt<br>ID) |
| --- | --- | --- | --- | --- |
| 23S |  |  |  |  |
| 792 | 747 | 5-Methyluridine | RlmC | RlmCD<br>(A0A0U1MPM5) |
| 1942 | 1915 | N5-Methylpseudouridine | RluD and<br>RlmH | RluD (Q2FX95)<br>and RlmH<br>(Q2G252) |
| 1966 | 1939 | 5-Methyluridine | RlmD | RlmCD<br>(A0A0U1MPM5) |
| 2278 | 2251 | 2'O-Methylguanosine | RlmB | RlmB (Q2G2M3) |
| 2472 | 2445 | N2-Methylguanosine | RlmKL | RlmK<br>(A0A0U1MLF0) |
| 2476 | 2449 | 5,6-Dihydrouridine | Unknown | - |
| 2530 | 2503 | 2'-Methyladenosine | RlmN | RlmN (Q2FZ66) |
| 16S |  |  |  |  |
| 535 | 527 | N7-Methylguanosine | RsmG | RsmG (Q2FUQ4) |
| 975 | 966 | N2-Methylguanosine | RsmD | RsmD (W8TQE6) |
| 976 | 967 | 5-Methylcytidine | RsmB | RsmB (Q2FZ67) |
| 1412 | 1402 | N4, O 2'-Dimethylcytidine | RsmI and<br>RsmH | RsmI (Q2G1S1)<br>and RsmH<br>(P60393) |
| 1509 | 1498 | 5-Methyluridine | RsmE | RsmE (Q2FXZ5) |
| 1529 | 1518 | N6-Dimethyladenosine | RsmA | RsmA (Q2G0T0) |
| 1530 | 1519 | N6-Dimethyladenosine | RsmA | RsmA (Q2G0T0) |

### Conclusions

In this work, we have determined high-resolution cryo-EM structures of FA and its derivative FA-CP in complex with the clinical drug target, the *S. aureus* ribosome-EF-G complex. The structures show two different states, a predominant CHI state and a less abundant POST state. In line with other recent cryo-EM studies, our structures suggest that the POST state of EF-G may only be stable in presence of FA, whereas EF-G, during unperturbed translocation, would quickly dissociate from the ribosome.

The first structures of FA-inhibited ribosome complexes were at resolutions where modelling of FA and its interactions with EF-G was challenging (16, 18, 19). Using the advances in cryo-EM, our high-resolution reconstructions allow accurate modeling of the FA molecule and its binding site in the clinical target. Moreover, we show that FA-CP binds in an identical position as FA and forms the same interactions with EF-G. In addition, the cyclopentane moiety of FA-CP has closer contacts with two side chains of EF-G and the SRL of 23S rRNA. There is additional space in this part of the binding pocket, which could be explored in future FA derivatives. The observed interactions suggest that the improved resistance profile is due to a higher binding affinity.

The structure also demonstrates how the higher-affinity derivative partially compensates for resistance mutations that weaken the direct FA-EF-G interactions or destabilize domain III and its interactions with domains I and II. Domain III of EF-G appears to have evolved to allow dynamics during the elongation cycle (38), but its stability is also key to FA inhibition. The high-resolution maps allowed building and refinement of high-quality models including many of the native rRNA modifications. The structure provides an accurate starting point for structure-guided drug discovery, not only of FA derivatives and analogues, but also of other antibiotics that target the *S. aureus* ribosome.

## MATERIALS AND METHODS

### Cloning, overexpression, and purification of *Staphylococcus aureus* EF-G

*S. aureus* EF-G *fusA* gene was amplified using Phusion master mix (Thermo Fisher Scientific, Waltham, MA, USA) according to the manufacturer’s instructions from a previous EF-G construct in pET30 (23) with the primers (5’)ATG GCT AGA GAA TTC TCA TTA GAA AAA ACT C and (5’)GTT ATT CAC CTT TAT TTT TCT TGA TAA TAT CTT CAG C and cloned into pEXP5-NT following the manufacturer’s instructions (pEXP5-NT_SaEF-G). The sequence was confirmed by sequencing. This construct encodes a 6xHis tag with a TEV cleavage site and a linker to the EF-G sequence. After TEV cleavage, only an additional Ser-Leu remains on the N-terminus.

pEXP5-NT_SaEF-G was transformed into *Escherichia coli* BL21 (DE3). An overnight culture was inoculated 1:100 into a 2.8L baffled flask with 800 ml of LB with 100 µg/ml of ampicillin. Protein expression was induced at an OD_600 nm_ of 0.5-0.6 with 1 mM isopropyl-β-thiogalactopyranoside and the culture was incubated for 16-18 h at 16 °C. The cells were harvested at 6000 rpm for 30 min in a JLA 9.1000 rotor (Beckman Coulter, Brea, California, USA), washed with 150 mM NaCl, and stored at −20 °C.

The cells were resuspended in lysis buffer (50 mM Tris-HCl pH 7.5, 300 mM NaCl, 5 mM β-mercaptoethanol) with 0.1 % (v/v) Triton X-100, one EDTA-free mini-Complete protease inhibitor tablet (Roche, Basel, Switzerland) and DNase I. Then, they were lysed in a flow cell disruptor (Constant Systems Ltd., Daventry, United Kingdom). The lysate was centrifuged at 16000 rpm for 45 min in an SS-34 rotor (Thermo Fisher Scientific).

The supernatant was filtered by 0.45 µm with a polyethersulfone syringe filter (Sarstedt AG & Co, Nûmbrecht, Germany) and incubated with 2 ml of Ni Sepharose Fast Flow (Cytiva, Uppsala, Sweden) equilibrated with lysis buffer. The column was washed with 25 column volumes of 10 mM imidazole in lysis buffer and the protein was eluted with 500 mM imidazole in lysis buffer. The eluate was exchanged back to lysis buffer using a PD-10 column (Cytiva, Uppsala, Sweden). The 6x-histidine-tag was removed by cleavage with 1:25 molar ratio of TEV protease at 8 °C for 16-18 h followed by reverse IMAC on 1 ml of Ni Sepharose Fast Flow. *S. aureus* EF-G was purified further by gel filtration on a Hiload 16/60 Superdex-200 column (GE Healthcare, Uppsala, Sweden) equilibrated with lysis buffer. The peak fractions were concentrated to 5-10 mg/ml with a 30-KDa cutoff Vivaspin Turbo 15 (Sartorius AG, Göttingen, Germany), flash-frozen in liquid nitrogen, and stored at −80 °C.

### Ribosome purification

200 ml of *S. aureus* NCTC 8325-4 culture was grown at 37°C with shaking (200 rpm) in Mueller Hinton Broth (MHB) and harvested in early logarithmic phase (A600 = 1.0 AU/ml). The cells were pelleted by centrifugation at 5000 x g, and weighed approximately 0.5 g. The cell pellet was resuspended in 50 ml of 20 mM Tris-HCl pH 7.5, 100 mM NH_4_Cl, 10 mM magnesium acetate, 0.5 mM EDTA, 3 mM β-mercaptoethanol (BME) and 0.3 μg/ml DNase I supplemented with 600 μl of protease inhibitor cocktail (Roche, Basel, Switzerland) and 5 mg lysostaphin (Sigma-Aldrich, Merck, Darmstadt, Germany) and incubated for 1 h at 37°C. The cells were then lysed by French press and the cell debris was pelleted and discarded by two centrifugations of 30 min at 20,000 rpm using a SS-34 rotor at 4°C. The supernatant was applied with volume ratio 1:1 on a 1.1M sucrose cushion in 20 mM Tris-HCl pH 7.5, 0.5 M NH_4_Cl, 10 mM magnesium acetate, 0.5 mM EDTA, 3 mM BME and centrifuged for 18 h at 30,000 rpm in a Ti 50.2 rotor (Beckman Coulter) at 4°C. The ribosome pellets were washed and dissolved by gentle stirring in the same buffer without sucrose and subjected to a second sucrose cushion centrifugation as above. Ribosome pellets were resuspended in buffer X (20 mM Tris-HCl pH 7.5, 60 mM NH_4_Cl, 5 mM magnesium acetate, 0.25 mM EDTA, 3 mM BME), and the 70S ribosomes were isolated by zonal centrifugation in a Ti 15 rotor (Beckman Coulter) for 15 h at 28,000 rpm on a sucrose gradient from 10 % to 40 % in buffer X. The 70S peak was collected, and the ribosomes were pelleted by ultracentrifugation in a Ti 50.2 rotor (Beckman Coulter) for 19 h at 38,000 rpm, resuspended in HEPES polymix buffer (5 mM HEPES pH 7.5, 5 mM NH_4_Cl, 5 mM Mg(OAc)_2_, 100 mM KCl, 0.5 mM CaCl_2_, 8 mM putrescine, 1 mM spermidine, and 1 mM dithioerythritol) (39), shock-frozen in liquid nitrogen and stored at −80°C.

### Complex preparation

The FA sample was prepared by mixing 0.5 µM (final concentrations) 70S *S. aureus* ribosomes in HEPES polymix buffer (with 20 mM HEPES pH 7.5 and 5 mM BME as reducing agent) with 2 µM mRNA (5’-GGCAAGGAGGUAAAAAUGGCAAAA-3’) and incubated for 10 min at 37 °C. Then, 2 µM *E. coli* tRNA^fMet^ was added and the mix was incubated for 10 min at 37°C. Next, 5 µM *S. aureus* EF-G, 400 µM FA (Sigma-Aldrich, Merck, Darmstadt, Germany), and 1 mM GTP were added and incubated for 10 min at 37°C, diluted 1:2, and kept on ice until plunge-freezing. The FA-CP sample was prepared as above, but with 5 µM *E. coli* tRNA^fMet^, and 400 µM FA-CP (28) instead of FA.

### Cryo-EM grid preparation and data collection

#### FA sample

A QuantiFoil 300-mesh R 2/2 grid with 2 nm continuous carbon (QuantiFoil Micro Tools GmbH, Großlöbichau, Germany) was glow-discharged 30 s at 20 mA and 0.39 mBar using an EasiGlow (Ted Pella, Inc., Redding, CA, USA). 2.5 µl of FA sample was incubated on the grid for 60 s. Then 1 µl of FusB 25 µM in polymix buffer with 400 µM FA was mixed into the drop, blotted for 4 s, and plunge-frozen in a Vitrobot mark IV (Thermo Fisher Scientific) at 4 °C and 95 % humidity. The grid was screened at the Uppsala Cryo-EM Facility, Sweden on a Glacios TEM operated at 200 kV equipped with a Falcon-III direct electron detector (Thermo Fisher Scientific). The final data was collected at SciLifeLab in Solna, Sweden on a Titan Krios G2 (Thermo Fisher Scientific) operated at 300 kV and equipped with a K3 BioQuantum direct electron detector (Gatan, Inc, AMETEK, Berwyn, PA, USA) and energy filter using 20 eV slit. The data was acquired at 105,000 x nominal magnification with a calibrated pixel size of 0.824 Å. A 50 µm C2 aperture was inserted and the TEM was operated in nanoprobe mode at spot size 6 and 670 nm beam size. The movies were collected in 30 frames over 1.3 s with a total dose of 30.83 e^-^/Å^2^ (16.1 e^-^/pixel/s) with a set defocus between −0.5 to −1.5 µm.

#### FA-CP sample

A Quantifoil 200-mesh R 2/1 grid with 2 nm continuous carbon was glow-discharged 15 s at 20 mA and 0.39 mBar using an EasiGlow. 3 µl of FA-CP sample was incubated on the grid for 30 s, then blotted for 4 s, and plunge-frozen in a Vitrobot mark IV at 4 °C and 95 % humidity. The grid was screened at the Uppsala Cryo-EM Facility as described above and the final data was collected at SciLifeLab in Umeå, Sweden in a Titan Krios G2 operated at 300 kV and equipped with a Falcon-4i direct electron detector and Selectris energy filter (Thermo Fisher Scientific) using 10 eV slit. The data was acquired at 165,000 x nominal magnification with a calibrated pixel size of 0.728 Å. A 50 µm C2 aperture was inserted and the TEM was operated in nanoprobe mode at spot size 6 and 520 nm beam size. The EER formatted movies were collected in 684 raw frames with a total dose of 27.77 e^-^/Å^2^ (12.45 e^-^/pixel/s) over 2.23 s with a set defocus between −0.7 to −1.3 µm.

### Cryo-EM data processing

#### FA dataset

The dataset was processed with cryoSPARC v3.3.1 (40) following the workflow in Figure S5 resulting in a 2.49 Å reconstruction of the FA-complex in the chimeric hybrid state.

#### FA-CP dataset

The dataset was processed with cryoSPARC v4.0.2 and v4.1.2 following the workflow in Figure S6 resulting in a 2.02 Å reconstruction of the FA-CP-complex in chimeric hybrid state and a 2.44 Å reconstruction of the FA-CP-complex in classical state. The CTFs were estimated using CTFFIND4 (41).

The FSC was calculated using cryoSPARC with a mask that was generated by low-pass filtering each map to 20 Å, thresholding to a level that covers the whole complex (0.1 for FA CHI complex, 0.06 for FA-CP CHI complex, and 0.07 for FA-CP POST complex), dilated 4 pixels and soft-padded 8 pixels. The final maps were post-processed using local filtering on cryoSPARC v4.2.1.

### Model building

Model building was started by individually rigid body fitting all chains from the 3.1 Å *S. aureus* 70S ribosome structure (PDBID: 7NHM) in ChimeraX v1.5 (42), excluding the tRNA into the FA-CP CHI state unsharpened map. 16S RNA was fitted in three different fragments (1-921, 922-1396, and 1397-1552). EF-G was rigid body fitted from the 1.9 Å x-ray crystal structure of *S. aureus* EF-G 2XEX in two different fragments (1-401 and 402-692). The E-site tRNA was rigid-body fitted from the 3.3 Å *E. coli* mid-translocation intermediate ribosome structure (PDBID: 7SSD). All chains were manually inspected and adjusted in Coot v0.9.8.5 (43), followed by real-space refinement in Phenix v1.20.1-4487-000 (44) against the local filtered map using secondary structure and Ramachandran restraints. This model was used as starting point for the FA CHI and the FA-CP POST structures and refined following the same procedure. For the FA-CP POST structure, the P-site tRNA was rigid body fitted from the 2 Å *E. coli* ribosome structure (PDBID: 7K00). Water molecules were added to the FA-CP structures using Coot. Coordination waters for magnesium ions were manually corrected. Model validation was performed using MolProbity (45) and problematic areas were inspected and corrected. The B-factors were refined against the unsharpened maps using Phenix. All ligand restraints were produced using the Grade web server (46) and were manually inspected and adjusted in all structures. Hidden surface area calculations were made using areaimol from CCP4 (47, 48). The structure figures were rendered using ChimeraX.

## Data availability

The cryo-EM maps and models in this study have been deposited in the Electron Microscopy Data Bank under the accession codes EMD-17365 (2.5 Å FA-complex with chimeric hybrid pe/E tRNA), EMD-17363 (2.4 Å FA-CP post-translocational state complex), and EMD-17364 (2 Å FA-CP-complex with chimeric hybrid pe/E tRNA), and Protein Data Bank under the accession codes 8P2H (2.5 Å FA-complex with chimeric hybrid pe/E tRNA), 8P2F (2.4 Å FA-CP post-translocational state complex), and 8P2G (2 Å FA-CP-complex with chimeric hybrid pe/E tRNA). Python custom scripts used during analysis and refinement are available on Git Hub (https://github.com/adriangzlz97/pdb_python_tools).

## Author contributions

A.G.L. and M.S. conceived the project; A.G.L. cloned and purified EF-G; R.K.K. and S.S. purified the *S. aureus* ribosomes; B.N.C. and P.J.H. synthesized FA-CP; A.G.L. prepared the cryo-EM sample; A.G.L. and D.L. screened and collected the cryo-EM data; A.G.L. processed the cryo-EM data and modeled the structures; D.L. and M.S. supervised the structural work; M.S secured funding; A.G.L. generated the figures; and A.G.L. and M.S. wrote the manuscript. All authors contributed to the final version of the manuscript.

## ACKNOWLEDGEMENTS

We thank Diarmaid Hughes for providing the *S. aureus* strain, Michael Hall for assistance with cryo-EM data collection, and Filipe Maia for maintaining our local computer cluster. We acknowledge the use of the Cryo-EM Uppsala facility for grid preparation and screening, funded by the Department of Cell and Molecular Biology, the Disciplinary Domains of Science and Technology and of Medicine and Pharmacy at Uppsala University. Cryo-EM data were collected at the Cryo-EM Swedish National Facility funded by the Knut and Alice Wallenberg Foundation, the Erling-Persson Family Foundation, and the Kempe Foundation; SciLifeLab; Stockholm University; and Umeå University.

This research was funded by grants from Uppsala Antibiotic Center to M.S., from the Swedish Research Council (2017-03827 and 2022-04511 to M.S.; 2018-05946 and 2018-05498 to S.S.; 2016-06264 to M.S. and S.S.) and from the Knut and Alice Wallenberg Foundation (KAW 2017.0055) to S.S.. A.G.L. has received a fellowship from the Sven and Lilly Lawski Foundation.

